# GenomicLinks: Deep learning predictions of 3D chromatin loops in the maize genome

**DOI:** 10.1101/2024.05.06.592633

**Authors:** Luca Schlegel, Rohan Bhardwaj, Yadollah Shahryary, Defne Demirtürk, Alexandre P. Marand, Robert J. Schmitz, Frank Johannes

## Abstract

Gene regulation in eukaryotes is partly shaped by the 3D organization of chro]matin within the cell nucleus. Distal interactions between *cis*-regulatory elements and their target genes are widespread and many causal loci underlying heritable agricultural traits have been mapped to distal non-coding elements. The biology underlying chromatin loop formation in plants is poorly understood. Dissecting the sequence features that mediate distal interactions is an important step toward identifying putative molecular mechanisms. Here, we trained GenomicLinks, a deep learning model, to identify DNA sequence features predictive of 3D chromatin interactions in maize. We found that the presence of binding motifs of specific Transcription Factor classes, especially bHLH, are predictive of chromatin interaction specificities. Using an *in silico* mutagenesis approach we show the removal of these motifs from loop anchors leads to reduced interaction probabilities. We were able to validate these predictions with single-cell co-accessibility data from different maize genotypes that harbor natural substitutions in these TF binding motifs. GenomicLinks is currently implemented as an open-source web tool, which should facilitate its wider use in the plant research community.

---

The spatial configuration of chromatin within the nucleus of eukaryotic cells is fundamental to genome regulation. Its structural dynamics are critical for various cellular processes, including DNA replication, repair, spatio-temporal gene expression patterning and transposable element silencing (1–3). Chromosome conformation capture (3C) coupled with next-generation sequencing (e.g. Hi-C (4), Hi-ChIP (5), ChIA-PET (6), Capture-C (7), Capture Hi-C (8), 4C (9), 5C (10) or In Situ Hi-C (11)) have emerged as powerful tools to interrogate 3D chromatin biology in a high-throughput fashion. These methods have led to systematic insights into the hierarchical organization of chromatin, including the identification of A/B compartments (4), Topologically Associated Domains (TADs) (12) and chromatin loops in the form of enhancer-promoter interactions (5). In animals, TAD and chromatin loop formation is largely facilitated by the CTCF-cohesin complex, which binds DNA to physically clamp down chromatin and force loop extrusion (11). Although the general principles of chromatin organization are conserved, plants are distinct in that they lack CTCF proteins. Perhaps as a result, TAD structures display less defined bound-aries and are often indistinguishable from local A/B compartments, particularly in large and complex plant genomes (13). How plants initiate and stabilize 3D chromatin interactions remains poorly understood. Several studies point to an enrichment of particular transcription factor (TF) families at loop anchors (14), which has led to the hypothesis that TFs mediate loop formation through processes like dimerization (15, 16). Hence, the presence of specific TF binding motifs may be an important determinant of 3D chromatin interactions. Testing this hypothesis in a high-throughput manner is experimentally challenging. Moreover, additional DNA sequence features, beyond known TF binding motifs, may be important contributors to loop formation, but remain difficult to identify. Machine learning (ML) methods trained on 3C data in animals have emerged as powerful tools to dissect the DNA sequence grammar underlying chromatin biology. Deep Learning (DL) models, in particular, have demonstrated remarkable success in predicting 3D chromatin contacts directly from DNA sequences (17), with prediction accuracies in the range of 0.7 to 0.9 (17). The power of these models is that they can be combined with *in silico* mutagenesis, where arbitrary DNA sequence mutations are induced and evaluated for their impact on looping probabilities. This approach provides a framework to systematically screen genetic variants underlying agronomically important complex traits that have been mapped to (distal) *cis*-regulatory elements (18). Examples of such traits in maize include *tb1* (19), *ZmCCT9, Vgt1, prol1*.*1*, and *UPA2* (20, 21). Such knowledge can generate concrete hypotheses for experimental validation using genome editing and/or present novel breeding targets. Harnessing DL models for studying plant chromatin biology is a promising goal. However, models trained on animal data perform poorly when applied to plants, yielding prediction accuracies as low as 0.1% (see SI material). This observation points to fundamental differences in the type of DNA sequence features that predict chromatin looping, and suggests that DL models need to be trained directly on plant genomic data to optimize performance. Here we implement such an approach. Using maize (*Zea mays*) as an experimental system, we developed GenomicLinks, a DL model capable of predicting 3D chromatin interactions from DNA sequences with high accuracy. Leveraging high-quality Hi-ChIP-seq dataset for training, we integrated the most successful architectural features from DL models used in the animal field. The model combines dual convolutional neural networks (CNNs) with a long short-term memory (LSTM) network, enabling identification of spatial and sequential aspects of chromatin interactions. Application of GenomicLinks coupled with a TF-centered *in silico* mutagenesis approach revealed binding motifs of specific Transcription Factor (TF) classes as crucial predictors of chromatin looping. We were able to validate these predictions with single-cell co-accessibility data from 9 different maize genotypes that harbor natural mutations in these TF binding motifs. GenomicLinks is currently implemented as an open-source web tool, which should facilitate its wider use among plant researchers.

## Material and Methods

### Hi-ChIP data collection

To construct a high-quality dataset of true chromatin interactions, we obtained Hi-ChIP data for the histone modifications H3K4me3 (associated with transcriptional activation) and H3K27me3 (associated with transcriptional repression) from Ricci et al. (2019) (18). These datasets were generated through an initial Hi-C process, followed by Chromatin Immunoprecipitation (ChIP), using antibodies against H3K4me3 (indicative of transcriptional activity) and H3K27me3 (indicative of transcriptional suppression), while excluding regions characteristic of constitutive heterochromatin (22–24). The Hi-ChIP raw data were processed using the HiC-pro pipeline (version 2.8.054) (25) and mapped to *Zea mays* B73 reference genome (version 4) (26), where alignments with a MAPQ score greater than five were retained. ChIP-seq pulldown efficiency was assessed through the analysis of dangling-end and self-ligation read pairs. Loop detection was performed using FitHiChIP (27) with valid read pairs, applying a 5 kb bin size, adjusting for coverage bias, and setting a false discovery rate (FDR) threshold of less than 0.01 to distinguish H3K4me3 and H3K27me3 Hi-ChIP loops. The sequenced data was also filtered to include only interactions with a minimum genomic distance of 20 kb and a maximum genomic distance of 2 Mb.

### Data preprocessing

To enhance accuracy and ensure our model’s comparability with similar deep learning frameworks (17, 28), we refined FitHiChIP’s original 5 kb anchor definitions to 2.5 kb regions centered on each anchor’s midpoint, to improve computational efficiency. Nucleotide sequences for these 2.5 kb windows were extracted from the *Zea mays* B73 reference genome (version 4) (26) for deep learning analysis. These sequences, encompassing 101,172 anchor pairs, constituted our set of true positive interactions and were labeled as ‘1’. To create the negative training set, we started with the positive set and permuted the anchor positions, ensuring that the negative set covered the same genomic space as the positive set for biological relevance. We kept a similar distance distribution of negative and positive samples. To improve the model’s ability to distinguish between positive and negative sets, we further adjusted the positions of each permuted anchor pair by randomly shifting it 5 to 25 base pairs left or right and labeled them as ‘0’. We found that permuting anchor pairs without these additional bp shifts result in no learning progress. This suggests that the model learns sequence features that determine the potential of two genomic regions to interact, rather than the 3D interaction architecture itself. Some alternative methods in the literature (29) opted to define negative sets by extracting random anchor sequences from the reference genome. In heterochromatin-rich genomes, like that of maize, this latter strategy runs into the danger that the models learn DNA features that distinguish euchromatin from heterochromatin, rather than interacting from non-interaction regions, and was therefore not employed here. Next, we combined our positive and negative training sets to form the “complete training set” (N=202,344). This complete set was shuffled to ensure that the order of the anchor pairs does not bias the model during training. Finally, one-hot encoding was applied to the 2.5 kb anchor pairs in the complete training set to convert the A, T, C, G nucleotide sequence into a binary matrix representation. These binary matrices along with their labels were stored in the H5 file format, making it compatible with TensorFlow (30) and Keras (31) for data handling during training.

### Model architecture and training process

We developed a dual CNN architecture to independently analyze each anchor, followed by an LSTM layer. This configuration has proven successful in animal systems (32),(33),(34), (35),(36). Our model takes two 2.5kb input anchor sequences as input, representing potential genomic interaction sites. The architecture employs a one-dimensional CNN with two convolutional layers, featuring filter sizes of 256 and 512, and kernel sizes of 64 and 32. These layers extract high-level features from the input sequences (37), which are then processed by max-pooling layers for dimensionality reduction. Subsequently, the outputs were concatenated and fed into one bidirectional LSTM layer, comprising 256 units. This LSTM layer effectively captures long-range dependencies within the input sequences (38). The final prediction was obtained through a dense layer with 512 units and a sigmoid activation function. To enhance generalization and prevent overfitting, we incorporated batch normalization and dropout layers. The optimization process utilizes the Adam optimizer with a learning rate of 1e-4, guided by the binary cross-entropy loss function. The entire architecture was implemented using the TensorFlow (30) and Keras (31) libraries (jlab github). The model summary, including detailed parameter configurations, is provided in supplementary materials (SI materials). During the training process, we continually monitored various statistical metrics. One crucial metric is specificity at a given sensitivity threshold for the validation set. This metric measures the model’s ability to correctly classify negative samples while maintaining a specified level of sensitivity. We utilized the EarlyStopping callback with patience of 10 epochs to ensure training halts when performance plateaus or deteriorates. ModelCheckpoint was employed to save the model weights only if they outperform the previous best epoch. The training process spanned 100 epochs, with each epoch comprising a batch size of 100 samples. We used the Adam optimizer to minimize the binary cross-entropy loss, a common choice for binary classification tasks (39). Detailed training logs and access to saved model weights can be found in our GitHub repository (jlab github).

### *In silico* mutagenesis of TFBS

To enable automated, targeted *in silico* mutagenesis of TFBS within loop anchors, we introduce SNIPER, an extension of the GenomicLinks toolkit, designed to analyze sequence features influencing genomic interactions (jlab github, Fig. 1B). SNIPER provides a targeted and a genome-wide mode for integrating *in silico* mutations, deletions or motif replacements.

**Figure 1:**
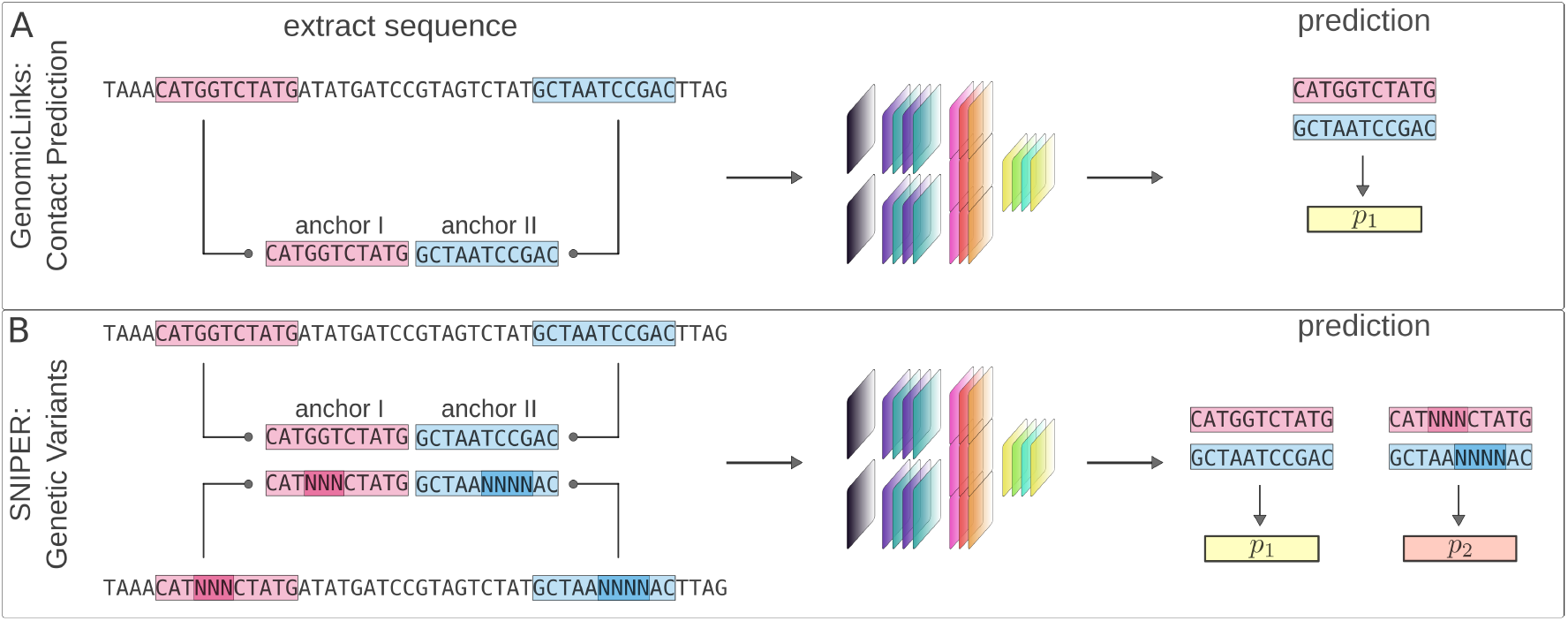
Workflow of GenomicLinks Deep Learning for predicting chromatin loops in the maize genome. **A)** GenomicLinks was trained on Hi-ChIP data to predict 3D chromatin interactions from DNA sequence features flanking chromatin loops. The trained model accepts any 2.5 kb anchor pairs as input and outputs an interaction probability *p*_1_. **B)** SNIPER is an *in silico* mutagenesis software extension embedded in the GenomicLinks framework. SNIPER can introduce targeted sequence mutations in selected anchor pairs and resupplies these to GenomicLinks for evaluation. GenomicLinks uses the mutated anchor pairs to assess chromatin loop stability, yielding interaction probabilities *p*_1_ for the original sequences and *p*_2_ for the mutated ones. This approach provides a setup to test the impact of specific genetic variants on chromatin loop stability.

### Web Server implementation

For web server implementation, a combination of Java scripting, PERL, HTML, CSS, and PHP was employed, all running on an Apache server utilizing the Hypertext Transfer Protocol (HTTP). The front-end of the web interface was written in JAVA scripts, PHP, and HyperText Markup Language (HTML). We utilized PHP and PERL for scripting common gateway interfaces and web-server interactions due to the platform independence and open-source nature of Apache, MySQL, and PHP technologies. A step-by-step user guide for GenomicLinks and SNIPER can be found in the SI materials.

### Evaluation of TFBS on chromatin looping using *in silico* mutagenesis

To determine the impact of transcription factor binding sites (TFBS) on Hi-ChIP loops, we used SNIPER to scan and replace nucleotides within any identified TFBS motifs, based on exact matches to the consensus sequence from JASPAR (40–46) database, with an ‘N’ to denote an unknown nucleotide. We assessed the impact of these modifications by comparing them with original predictions. Anchor pairs with a probability drop of more than 50% post-modification were classified as ‘Relevant Anchor Pairs’ (RAPs) and their associated motifs were termed ‘relevant motifs’. Focusing on chromosome 1 as a test case, we identified 670 RAPs, encompassing 25,287 unique sequences with an average motif length of 8.5 base pairs. For comparison, we created a set of Control Anchor Pairs based on control motifs, which had minimal impact on loop probability, under comparable conditions of count and motif length. Notably, these control motifs were more prevalent, occurring in 2,623 Control Anchor Pairs.

### Validation of *in silico* predictions using maize genetic variation data

We process single nucleotide variant (SNV) data from the maize 282 diversity panel (47). The complete SNV dataset comprised approximately 2.2 million single nucleotide polymorphisms (SNPs) or indels, exclusively for chromosome 1. To prepare the dataset for GenomicLinks analysis, we updated the Hi-ChIP anchor pairs for each of the 278 maize inbred lines. We extracted DNA sequences from the reference genome and incorporated mutations as specified by the SNV matrix. Mutations were updated accordingly: unknown nucleotides (‘N’) were encoded as a uniform distribution vector (0.25, 0.25, 0.25, 0.25)^T, while deletions (‘D’) were represented by a zero vector (0, 0, 0, 0)^T. It is important to note that insertions (‘I’) were excluded from this process. We applied GenomicLinks to predict chromatin interactions within chromosome 1 for all 278 samples, creating a matrix of dimensions 278 x 16,410.

### Validation of *in silico* predictions using maize single-cell co-accessibility data

To further validate the *in silico* predictions that specific SNVs affect chromatin loop probabilities, we analyzed single-cell ATAC-seq data (48) for 10 genotypes apart of the maize 282 diversity panel (B73 (reference), B97, CML103, CML52, KI3, Ky21, M162W, M37W, OH7B, and Oh43), and processed the sequenced outputs as previously described (49). To identify regions of co-accessibility as a proxy for chromatin interactions, we first partitioned the nuclei by accessible chromatin regions (ACRs) matrix to restrict the analysis to each genotype in isolation. Nuclei from a single genotype were then aggregated in *X* pseudocells using *k* nearest neighbors, where *X* is an integer determined as the number of nuclei from the focal genotype divided by *k*, and *k* is an integer of the square root of the total number of nuclei from the focal genotype. Tn5 integrations per pseudocell across all ACRs were then scaled per million using the function *cpm* from the R package, *edgeR (50)*. Initial values of co-accessibility between ACRs were estimated as Spearman Correlation Coefficients across pseudocells, conditioning ACRs to be less than 500-kb and more than 2-kb apart. We assessed whether co-accessibility values dropped in anchor pairs, whose SNVs in RAPs were predicted to result in a loss of interaction probability. As a control, we assessed decreases in co-accessibility values in RAPs, where SNVs did not lead to predicted losses in interaction probabilities.

## Results

### Description of GenomicLinks

GenomicLinks is an open-source web tool designed to predict 3D chromatin interactions from DNA sequence in maize. GenomicLinks builds on a deep learning model, comprising two CNNs and LSTMs, and has been trained on curated maize Hi-ChIP data (Figure 1 and Methods). Users can upload a pair (or multiple pairs) of genomic sequences in either FASTA or BEDPE formats. These inputs are automatically processed and submitted to the trained model for interaction predictions. GenomicLinks outputs interaction probabilities for each sequence pair from the BEDPE files or with each individual sequence pair from FASTA files (Figure 2).

**Figure 2:**
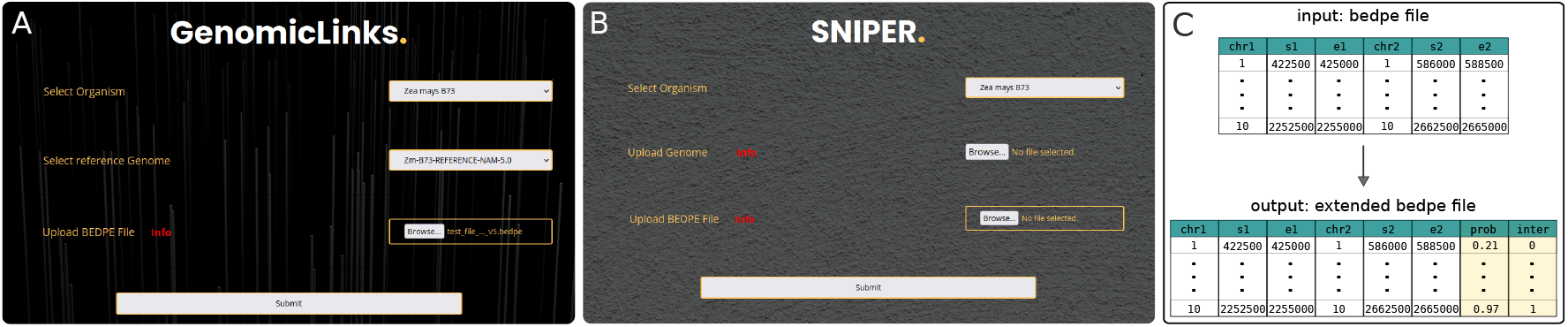
GenomicLinks user interface. **A)** The GenomicLinks web portal interface for uploading BEDPE files, currently supporting *Zea mays* with options for reference genome versions v4 or v5. **B)** Display of the input format required, and the output generated by GenomicLinks, adding predicted interaction probabilities (prob) and interaction presence (inter) for chromatin anchor pairs (*s*_1_, *e*_1_ and *s*_2_, *e*_2_). **C)** Display of the input format required and the output generated by GenomicLinks, adding predicted interaction probabilities (prob) and interaction presence (inter) for chromatin anchor pairs (*s*_1_, *e*_1_ and *s*_2_, *e*_2_).

### GenomicLinks displays favorable classification performance

We evaluated GenomicLinks’ classification performance using standard metrics (see Figure 3). The performance evaluation included the complete training data (details in the Materials and Methods), consisting of 101,172 true positive and an equal number of true negative interactions. This dataset was partitioned into training (80%, 161,875 samples), validation (10%, 20,234 samples), and test (10%, 20,234 samples) sets. Applying the trained model to the test set yielded favorable performance statistics: Accuracy (0.848), Sensitivity (Recall, 0.853), Specificity (0.843), F1-Score (0.848), Precision (0.844), ROC AUC (0.925), and AUPRC (0.926). Although these performance values were in line with previous deep learning approaches in animal genomes (32),(33),(34),(35),(36), GenomicLinks learned very different DNA sequence features predictive of chromatin looping. To illustrate this, we applied DeepMILO (32), a model originally designed for evaluating noncoding variant impacts on CTCF/cohesin-mediated insulator loops in the human genome, to the maize Hi-ChIP loops. DeepMILO predicted only 0.1% of Hi-ChIP loops correctly, corresponding to 144 out of 101,172 loops (SI material).

### *In silico* predictions of TF mediated 3D chromatin contacts

We set out to identify specific DNA sequence features underlying our DL predictions. Several studies have pointed to an enrichment of particular classes of TFs at loop anchors (15, 16). This has led to the hypothesis that TFs may mediate loop formation through processes like dimerization/protein interactions (51). To test whether the presence of TFBS in loop anchors contribute to our predictions, we applied SNIPER to perform a TF-centered *in silico* mutagenesis (see Materials and Methods). Focusing on chromosome 1 as a test case, we selected anchor pairs, whose 3D interactions were supported empirically by Hi-ChIP data as well as computationally by GenomicLinks (N = 16,083 out of 16,410). Utilizing the JASPER database, we identified all high-confidence TFBS for maize (82,000) within these sequences and applied SNIPER for targeted deletions of these sites (see Materials and Methods and Figure 1B). We observed that all anchor pairs contained at least one TFBS, with over 95% of these anchors having less than 20% of their base pairs overlapping TFBSs. The mutated sequences were then re-analyzed using GenomicLinks for interaction prediction. Our analysis identified a subset of approximately 4% of anchor pairs (678 out of 16,083) where TFBS deletions led to a strong probability shift of at least 50%, indicating a shift from interacting to non-interacting state. We refer to this subset as Relevant Anchor Pairs (RAPs). This results highlights cases where removal of only approximately 205 bp motif sequence from 5 kb loop anchors (on average) are sufficient to compromise looping probabilities. For comparison, we also included two benchmark scenarios into our analysis. In the first, we introduced random SNPs matching the total base pairs of all motifs within each anchor. In the second, we randomly repositioned deleted motif segments within the anchor sequence (see Figure 3C). No loss of interaction probabilities could be detected in the two control benchmark scenarios (Figure 3C), thus providing additional evidence that these motif sequences drive the predictions. Further examination of the frequency distribution of specific TF classes in the RAPs compared with all loops revealed significant shifts in class distribution (see Figure 3B). The most significant changes were found for the *AP2/EREBP* and *Basic helix-loop-helix* (*bHLH*) classes (*χ*^2^ = 87.21, df = 6, P = 1.85e-23), with the *AP2/EREBPs* showing a substantial depletion in the RAPs (all loops: 72.8%, RAPs: 52%), and the *bHLH* s an enrichment (all loops: 8.7%; RAPs: 18.7%). Interestingly, in mammals, members of the *bHLH* class have been shown to cooperate with histones to facilitate DNA access (52), and contribute to the 3D rewiring of chromatin architecture during cell lineage transitions (53).

**Figure 3:**
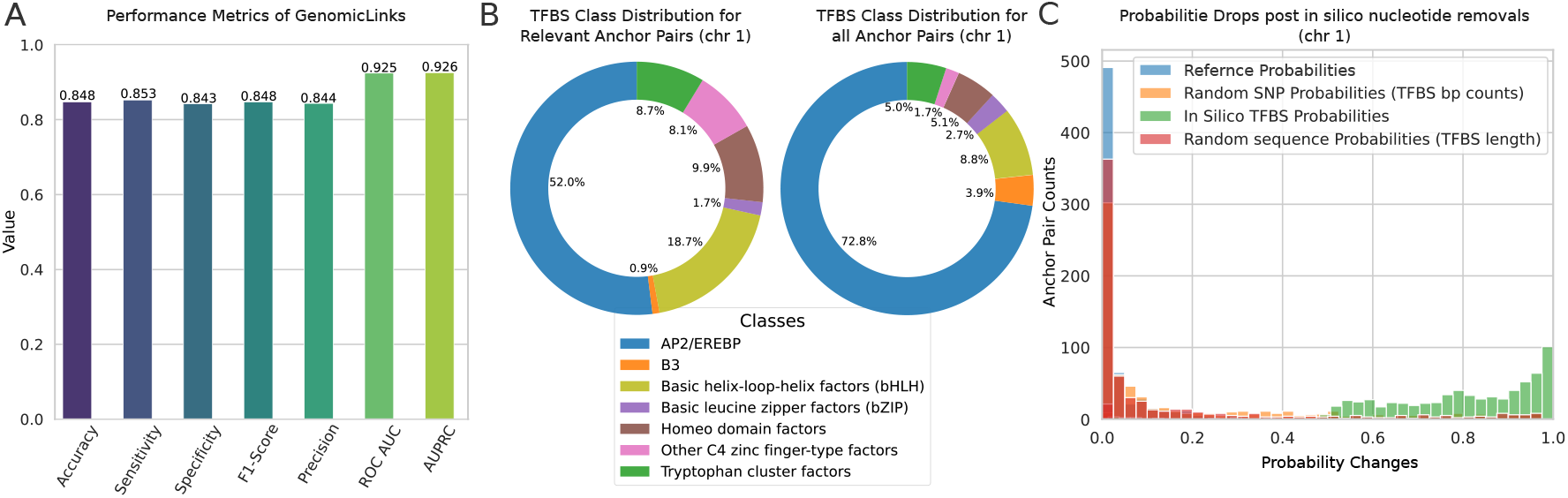
GenomicLinks performance statistics and impact analysis of *in silico* mutagenesis of Transcription Factor Binding Sites. **A)** Performance metrics, including accuracy, sensitivity, specificity, F1-score, precision, and AUC values. **B)** *Left:* Distribution of TFBS motif classes within all 16,410 Hi-ChIP anchor pairs on chromosome 1 of the *B73* reference genome. *Right:* Distribution TFBS motif classes within anchor pairs predicted to lose interaction after removal of TFBS, with *bHLH* (yellow) and *Other C4 zinc finger factors* (pink) significantly affecting loop stability. **C)** Histogram displaying the effect of *in silico* TFBS motif knockout on interaction probability: original (blue) versus collapsing interactions (green), alongside control scenarios (red/orange) verifying the specificity to the targeted TFBS motifs and their position.

### Natural mutations in TFBS lead to predicted losses in 3D chromatin contacts

The above analysis demonstrated that *in silico* deletions of entire TFBS can lead to a loss of predicted chromatin interactions in a subset of regions. In natural settings, complete deletions of TFBS are less common than single nucleotide variants (SNVs) in the form of SNPs or small indels. We asked if naturally occurring SNVs among maize inbred lines in these TFBS are sufficient to compromise predicted looping probabilities. To assess this, we screened genotype data from 9 maize NAM founder lines, and retained only those RAPs that contained at least one SNPs or small indels in a TFBS (Materials and Methods). Depending on the NAM line, this resulted in 172 to 551 RAPs (average of 487) on chromosome 1 for downstream analysis. Employing SNIPER (see Figure 1B), we replaced the B73 sequence in each TFBS with the SNP/indel genotype of the respective NAM line. On average, this translated to only 4.9 bp being substituted in a total of 205 bp of TFBS space per anchor pair (i.e. 2.39% of the TFBS space). After re-applying GenomicLinks to the SNV substituted RAPs, we found that 13% (average of 65 RAPs) exhibited a drop in predicted interaction probability of at least 10%, thus demonstrating considerable sensitivity to minor genetic variation (Figure 4C).

**Figure 4:**
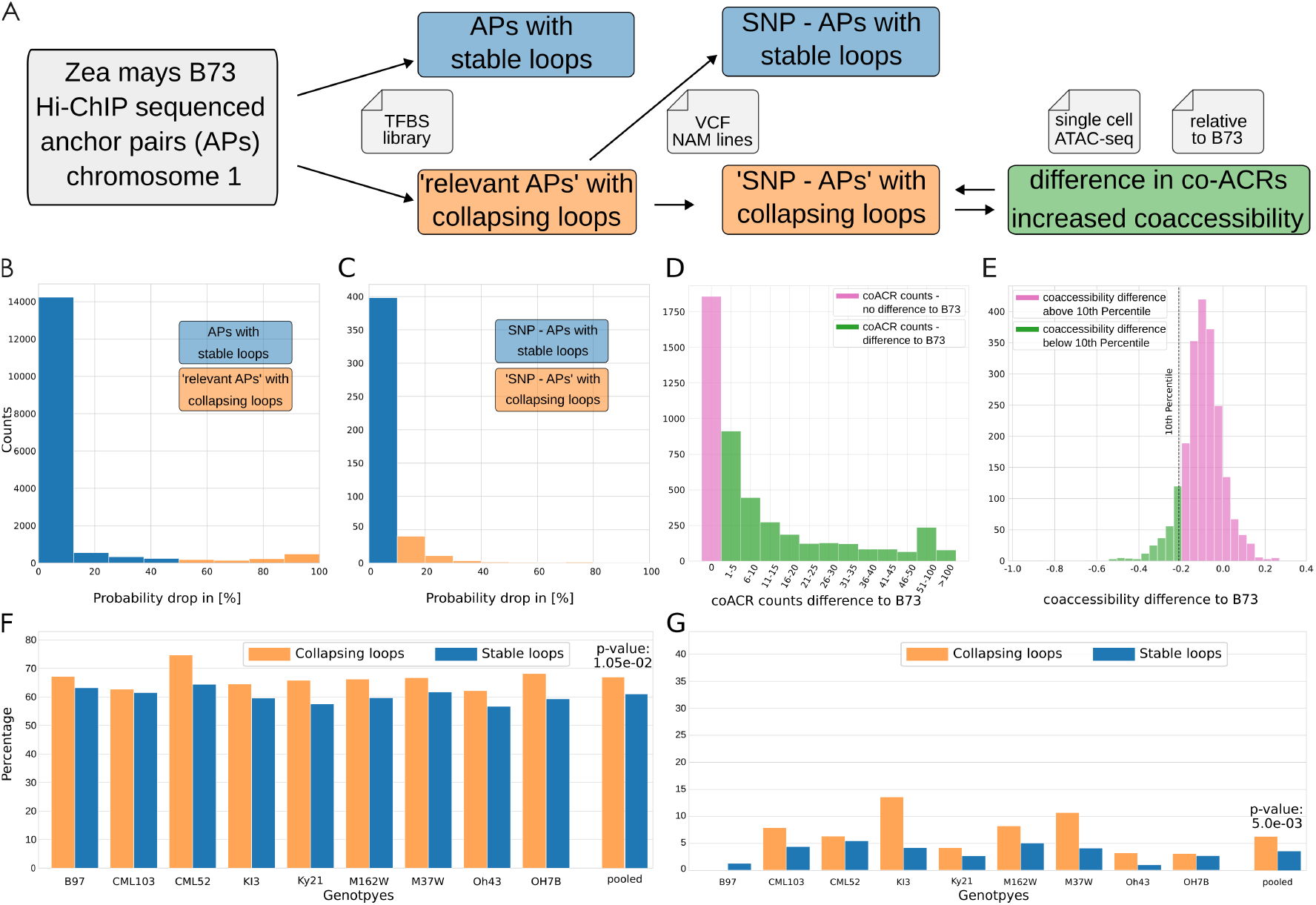
Validating GenomicLinks predictions using natural genetic variation and single cell ATAC-seq data. **A)** Workflow diagram illustrating the process for extracting relevant Anchor Pairs (APs) from Hi-ChIP sequenced data on chromosome 1 of *Zea mays B73*. APs are categorized based on their loop stability after *silico test* removal of Transcription Factor Binding Sites (TFBS) motifs. Further analysis mutates single nucleotide variants (SNVs) across 10 genotypes to these APs, we call them ‘SNP-APs’. Evaluation of ‘SNP-APs’ reveals a higher occurrence of collapsing loops in those with reduced coACR counts and increased co-accessibility differences relative to the B73 reference. **B)** Predicted probability drops of APs post-TFBS *silico test* removal indicate a majority of stable loops, defined by a drop of less than 50% from the original loop prediction for B73. **C)** Predicted probability drops of relevant APs post-SNV *silico test* mutation show a delineation between stable and collapsing loops, now with a tighter 10% cutoff from the original prediction for B73. **D)** Distribution of co-ACR count differences for all SNP-APs relative to *B73*, highlighting those with a positive difference (labeled green) for further loop analysis. **E)** Distribution of average co-accessibility differences for all SNP-APs compared to *B73*, highlighting those below the 10th percentile (labeled green) for further loop analysis. **F)** Normalized analysis for all genotypes reveals that collapsing loops more frequently occur than stable ones in SNP-APs with positive coACR count differences (green subset from figure D). **G)** Normalized analysis across all genotypes indicates that collapsing loops are more frequent than stable ones in SNP-APs with co-accessibility differences below the 10th percentile (green subset from figure E), though B97 deviates from this trend.

### Single-cell co-accessibility validates predicted interaction losses

To empirically validate our predicted interactions losses in the 487 RAPs, we analyzed single-cell ATAC-seq data for 10 NAM lines (48), and quantified chromatin co-accessibility, which has been widely used as a proxy for chromatin interactions (see Material and Methods) (54). As a control, we performed the same quantification in RAPs where SNP/indels mutations had no predicted effect on looping. We found that RAPs that exhibited reduced loop stability as a result of SNP/indel mutations within TFBS also showed a corresponding decrease in co-accessibility strength across cells compared to the B73, with B97 being the exception (Pooled genotype test, *χ*^2^ = 7.89, df = 1, P = 5.0e03). A similar trend was observed when comparing the number of co-accessible chromatin regions (co-ACR) instead co-accessibility strength (Pooled genotype test, Fisher’s Exact Test = 0.82, P = 1.05e-02), which indicated that TFBS-associated SNPs/indels also reduced the extent of open chromatin in the RAPs.

### Predicting the impact of genetic variation on chromatin looping in 277 maize inbred lines

We sought to extend our genotype-based predictions of differential chromatin looping to the panel of 277 maize diverse inbred lines (47), which has become an important genomic resource in maize. To that end, we identified all APs genome-wide containing published SNVs (101,172 APs genome-wide). This analysis yielded a large matrix of individual loop predictions (277 x 101,172). We identified significant variability in loop stability across anchor pairs among genotypes (SI Table 1, Fig. 5B). Interestingly, certain genotypes displayed a general trend towards loop loss (e.g. NC328, CI66 and A641), with an average loop stability of only 76% genome-wide. As a control, we examined a specific anchor pair on chromosome 1 surrounding the *tb1* locus, which is a well-characterized distal *cis*regulatory element known to form stable loops in modern maize (18, 19, 55). Indeed, despite the presence of genetic variation around this locus, with some genotypes showing base substitution in up to 454 bp within this anchor pair, loop probability remained consistently high (>99%) in all 277 inbred lines (Fig. 5A). Our complete computational predictions could be tested in future empirical studies of the diversity panel using a combination of targeted QTL mapping and population-level single-cell co-accessibility and/or HiC data. Overall, our results demonstrate that GenomicLinks learned meaningful DNA sequence features predictive of chromatin looping in maize, and that the model can be used to test the impact of specific genomic variants on chromatin looping.

**Figure 5:**
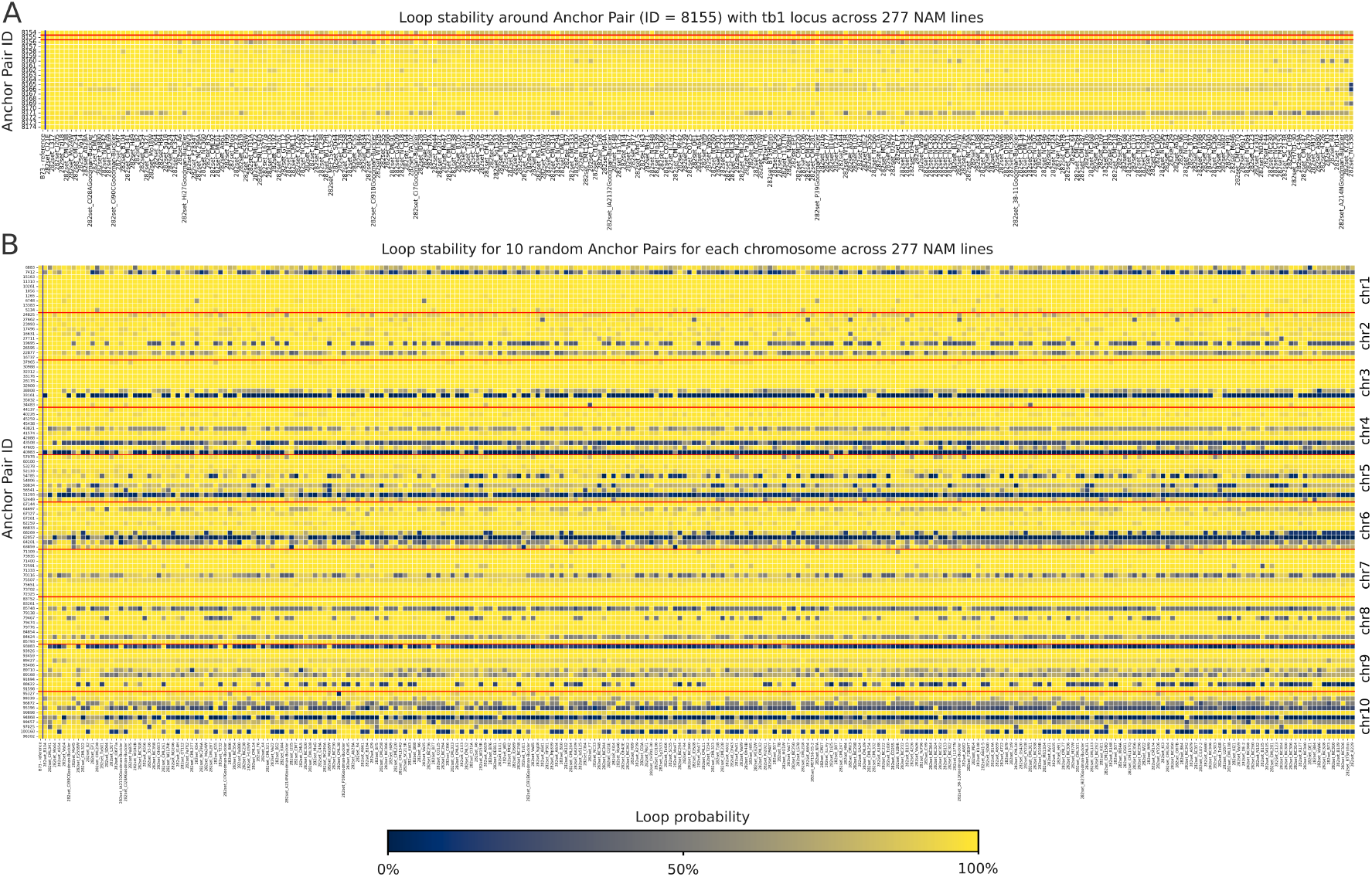
Impact of Genetic Variation on Chromatin Looping. **A)** Interaction probabilities for anchor pairs in the region chr1:270276250-270488750, including the tb1 locus, across 277 maize inbred lines, demonstrating consistent loop stability for tb1. **B)** Interaction probabilities visualized for 10 randomly selected anchor pairs across all 10 chromosomes, illustrating variability in chromatin loop stability. The analysis reveals genotypes with a higher tendency to lose loop stability (right), identifies regions that consistently exhibit stable interactions, and pinpoints anchor pairs prone to collapse across almost all genotypes.

## Discussion

Deep learning models like GenomicLinks have the potential to reveal complex and nested sequence motifs and genomic features responsible for the 3D structure of chromatin, which traditional sequence-based methods might miss. This computational framework enables an in-depth analysis of how genetic variants influence chromatin loop stability, a key factor in gene regulation. Additionally, it facilitates the exploration of regulatory variations associated with complex traits and investigates how non-coding loci, through physical interactions with distal genes, could modify gene expression patterns and influence phenotypic outcomes. Yet, GenomicLinks has several limitations that are inherent to the ‘black box’ nature of deep learning models. The model’s decision-making process is usually obscured by the complex interplay of millions of parameters across its node and layer architecture. The nested neural structure with its nonlinear operations between each layer makes it challenging to directly interpret how it evaluates and integrates local and distal genomic features. We already introduced a way to try to interpret DL decisions: *in silico* mutagenesis, where we strategically change sequences to observe how such changes impact model predictions. This method can be useful to identify crucial sequence features for the model’s decisions, labeling significant genomic regions for chromatin interaction predictions. In the context of deep learning analysis, this approach is known as input perturbation, which systematically tests the model’s sensitivity and reliance on specific input features. To further address the ‘black box’ nature of deep learning models, Class Activation Mapping (CAM) provides a more direct method for visualizing influential input segments. CAM produces heat maps that highlight which parts of the input significantly influence the model’s decisions, offering insights into the impactful regions within the input sequence for specific anchor pairs. However, CAM’s utility is primarily limited to the convolutional layers, and it does not adequately address the complexities introduced by LSTM layers, which are crucial for capturing long-distance dependencies within the chromatin. Therefore, while CAMs can be valuable for in-depth analysis of particular anchor pairs, they are less effective for comprehensive analysis across large datasets. Based on work in animals, it is unlikely that GenomicLinks can be directly applied to other plant species without re-training (56, 57). However, transfer learning is a promising strategy to mitigate this limitation. The concept of transfer learning involves retraining the model, initially trained on a specific species like *Zea mays*, on new datasets from different species. This process allows the model to adapt, learning both species-specific features and universal chromatin organization principles. Furthermore, it provides a method for estimating the proportion of chromatin features that are universal versus those that are genome-dependent by comparing models trained directly on various species and those adapted via transfer learning.

## Acknowledgements

This work was partly supported by the Technical University of Munich Institute for Advanced Study, funded by the German Excellence Initiative and the European Seventh Framework Programme under grant agreement no. 291763. We thank members of the Johannes lab for helpful feedback. This research was supported by the National Science Foundation (IOS-1856627) and the UGA Office of Research to RJS, and the National Institute of General Medical Sciences of the National Institutes of Health (1R00GM144742) to APM.

