## Supplemental Information for "GenomicLinks: Deep learning predictions of 3D chromatin loops in the maize genome"

### Supplemental Information - GenomicLinks

**Performance of human-trained models for Zea mays HiChIP sequenced loops**

We adapted the human-genome-based deep learning model, DeepMilo (1), initially trained on the hg19 reference genome (2), to analyze chromatin interactions in maize. To align the model with our species of interest, we substituted the human reference genome with the Zea mays genome version 4 (3). In a preliminary assessment without integrating mutations, we observed that the model failed to accurately predict over 99.9% of the maize chromatin loops. This approach highlights the need for species-specific or condition-specific model training to achieve reliable predictive performance in genomic studies. The prediction results are provided in https://github.com/jlab-code/GenomicLinks.

```
Model: "GenomicLinks"
__________________________________________________________________________________________________
 Layer (type)                   Output Shape         Param #     Connected to
==================================================================================================
 input_3 (InputLayer)           [(None, 2500, 4)]    0           []

 input_4 (InputLayer)           [(None, 2500, 4)]    0           []

 conv1d_4 (Conv1D)              (None, 2437, 256)    65792       ['input_3[0][0]']

 conv1d_6 (Conv1D)              (None, 2437, 256)    65792       ['input_4[0][0]']

 max_pooling1d_4 (MaxPooling1D)  (None, 1217, 256)   0           ['conv1d_4[0][0]']

 max_pooling1d_6 (MaxPooling1D)  (None, 1217, 256)   0           ['conv1d_6[0][0]']

 conv1d_5 (Conv1D)              (None, 1186, 512)    4194816     ['max_pooling1d_4[0][0]']

 conv1d_7 (Conv1D)              (None, 1186, 512)    4194816     ['max_pooling1d_6[0][0]']

 max_pooling1d_5 (MaxPooling1D)  (None, 592, 512)    0           ['conv1d_5[0][0]']

 max_pooling1d_7 (MaxPooling1D)  (None, 592, 512)    0           ['conv1d_7[0][0]']

 concatenate_1 (Concatenate)    (None, 1184, 512)    0           ['max_pooling1d_5[0][0]',
                                                                  'max_pooling1d_7[0][0]']

 dropout_2 (Dropout)            (None, 1184, 512)    0           ['concatenate_1[0][0]']

 bidirectional_1 (Bidirectional  (None, 512)         1576960     ['dropout_2[0][0]']
 )

 flatten_1 (Flatten)            (None, 512)          0           ['bidirectional_1[0][0]']

 dense_2 (Dense)                (None, 512)          262656      ['flatten_1[0][0]']

 batch_normalization_1 (BatchNo  (None, 512)         2048        ['dense_2[0][0]']
 rmalization)

 dropout_3 (Dropout)            (None, 512)          0           ['batch_normalization_1[0][0]']

 dense_3 (Dense)                (None, 1)            513         ['dropout_3[0][0]']

==================================================================================================
Total params: 10,363,393
Trainable params: 10,362,369
Non-trainable params: 1,024
__________________________________________________________________________________________________
```

**Figure 1: GenomicLinks - model and parameter overview model.summary()**

The model summary outlines the architecture of the GenomicLinks deep learning model, detailing each layer's configuration, including the number of parameters in convolutional, pooling, LSTM, and dense layers. It has been trained on a Nvidia GTX 3090 TI, with each epoch completing in less than 10 minutes. All scripts and data which have been used can be found in https://github.com/jlab-code/GenomicLinks.

**User Input Navigation for GenomicLinks**

- Select Reference Genome:
  - Begin by choosing your reference genome. Currently, GenomicLinks supports Zea Mays B73 version 4 or version 5.
- Upload Your BEDPE File:
  - Upload a BEDPE file that defines your regions of interest. This file should contain at least six essential columns: chr1, start1, end1, chr2, start2, end2. These columns specify the chromosome and start/end positions of each anchor pair. Note that additional columns in the file will be ignored, and anchor size is set to a length of 2500 bp around the center for uniform analysis.
- Initiate Analysis and Access Your Results:
  - After selecting your reference genome and uploading your BEDPE file, click 'Submit' to start the analysis. GenomicLinks will then begin processing your data to predict chromatin interactions for the specified anchor pairs. This process may take several minutes, depending on the size of your input. Once completed, you will be provided with a link to access your results. The output includes interaction probabilities for each anchor pair, indicating the likelihood of interaction based on the genomic features and sequences.

**User Input Navigation for SNIPER**

- Prepare Your Genome Data:
  - Start by preparing your mutated reference genome in a FASTA format. Currently, SNIPER is optimized to process single chromosomes, with a file size limit up to 50MB.
- Define Regions of Interest:
  - Upload a BEDPE file to specify your regions of interest, just like in GenomicLinks. This file should list the chromosome and start/end positions for each anchor pair within six essential columns: chr1, start1, end1, chr2, start2, end2.
- Initiate Analysis and Access Your Results:
  - Once your mutated genome data and BEDPE file are prepared, click 'Submit' to start the analysis. SNIPER will extract regions from your specified genome and predict chromatin interactions for each anchor pair. Results will be ready for download shortly, the duration depends on the size of the input.
  - To evaluate the impact of mutations on loop stability, you can perform a comparative analysis by repeating the process with the original, non mutated genome. This comparison will highlight changes in interaction probabilities, providing insights into how specific mutations may influence chromatin loop stability.

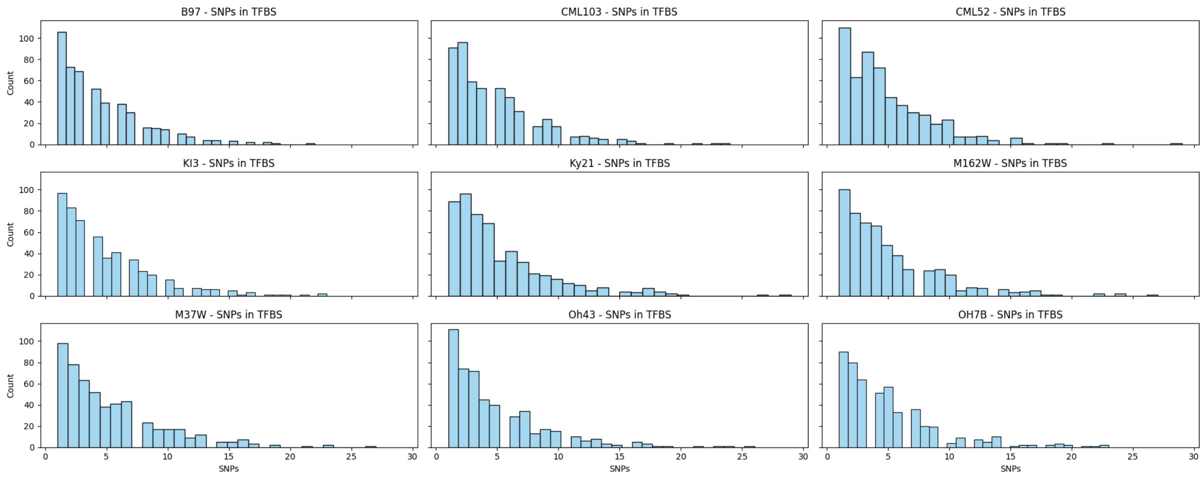

**Figure 2:** Distribution of SNPs in Transcription Factor Binding Sites (TFBS) across 9 genotypes.

| | chr1 | s1 | e1 | chr2 | s2 | e2 | B73 - reference | 282set_33-16 | 282set_38-11Goodman-Buckler | 282set_4226 | ... | 282set_VaW6 | 282set_W117Ht | 282set_W153R | 282set_W182B | 282set_W22 | 282set_W64A | 282set_WD | 282set_Wf9 | 282set_Yu796 | 282set_i1677a |
|---|---|---|---|---|---|---|---|---|---|---|---|---|---|---|---|---|---|---|---|---|---|
| **0** | 1 | 41250 | 43750 | 1 | 91250 | 93750 | 0.999776 | 0.999898 | 0.999895 | 0.999898 | ... | 0.999895 | 0.999764 | 0.999896 | 0.999765 | 0.999895 | 0.999895 | 0.999896 | 0.999895 | 0.999895 | 0.999895 |
| **1** | 1 | 227156250 | 227158750 | 1 | 227191250 | 227193750 | 0.996870 | 0.502070 | 0.381758 | 0.394144 | ... | 0.396570 | 0.393965 | 0.955635 | 0.876898 | 0.392785 | 0.392976 | 0.389714 | 0.112216 | 0.216308 | 0.394670 |
| **2** | 1 | 227156250 | 227158750 | 1 | 227221250 | 227223750 | 0.995450 | 0.754709 | 0.745049 | 0.746568 | ... | 0.754827 | 0.752091 | 0.653338 | 0.756005 | 0.777897 | 0.750153 | 0.738166 | 0.827414 | 0.680795 | 0.995541 |
| **3** | 1 | 227156250 | 227158750 | 1 | 227281250 | 227283750 | 0.999998 | 0.999043 | 0.999227 | 0.999251 | ... | 0.999252 | 0.999998 | 0.999259 | 0.999998 | 0.999833 | 0.999242 | 0.999825 | 0.999242 | 0.999645 | 0.999241 |
| **4** | 1 | 227156250 | 227158750 | 1 | 227371250 | 227373750 | 0.999953 | 0.996091 | 0.997715 | 0.568466 | ... | 0.189199 | 0.987211 | 0.987793 | 0.623641 | 0.634812 | 0.987482 | 0.627343 | 0.987482 | 0.702263 | 0.568081 |
| **5** | 1 | 227156250 | 227158750 | 1 | 227386250 | 227388750 | 0.999947 | 0.999944 | 0.999945 | 0.999943 | ... | 0.999946 | 0.999960 | 0.999947 | 0.999945 | 0.999947 | 0.999946 | 0.999946 | 0.999946 | 0.999948 | 0.999944 |
| **6** | 1 | 227156250 | 227158750 | 1 | 227426250 | 227428750 | 0.999999 | 0.999959 | 0.999963 | 0.999958 | ... | 0.999960 | 0.999966 | 0.999992 | 0.999961 | 0.999961 | 0.999963 | 0.999962 | 0.999964 | 0.999992 | 0.999965 |
| **7** | 1 | 227156250 | 227158750 | 1 | 227451250 | 227453750 | 0.999615 | 0.999605 | 0.998860 | 0.998278 | ... | 0.998264 | 0.998290 | 0.998827 | 0.998357 | 0.998283 | 0.998273 | 0.998315 | 0.998268 | 0.998283 | 0.998319 |
| **8** | 1 | 227166250 | 227168750 | 1 | 227216250 | 227218750 | 0.976803 | 0.004049 | 0.004014 | 0.004117 | ... | 0.004283 | 0.015030 | 0.294151 | 0.028364 | 0.004043 | 0.015290 | 0.004247 | 0.015108 | 0.011899 | 0.005822 |
| **9** | 1 | 227176250 | 227178750 | 1 | 227226250 | 227228750 | 0.995005 | 0.967778 | 0.974203 | 0.973886 | ... | 0.968300 | 0.993014 | 0.969224 | 0.997301 | 0.997193 | 0.992900 | 0.997257 | 0.992711 | 0.734826 | 0.952492 |

10 rows × 284 columns

| | chr1 | s1 | e1 | chr2 | s2 | e2 | B73 - reference | 282set_33-16 | 282set_38-11Goodman-Buckler | 282set_4226 | ... | 282set_VaW6 | 282set_W117Ht | 282set_W153R | 282set_W182B | 282set_W22 | 282set_W64A | 282set_WD | 282set_Wf9 | 282set_Yu796 | 282set_i1677a |
|---|---|---|---|---|---|---|---|---|---|---|---|---|---|---|---|---|---|---|---|---|---|
| **101162** | 10 | 72986250 | 72988750 | 10 | 73136250 | 73138750 | 0.999840 | 0.999939 | 0.999930 | 0.999840 | ... | 0.999973 | 0.999930 | 0.999983 | 0.999927 | 0.999854 | 0.999940 | 0.999843 | 0.999973 | 0.999837 | 0.999845 |
| **101163** | 10 | 72986250 | 72988750 | 10 | 73056250 | 73058750 | 0.999477 | 0.999475 | 0.999756 | 0.999447 | ... | 0.999741 | 0.999749 | 0.999753 | 0.999754 | 0.999741 | 0.999467 | 0.999476 | 0.999741 | 0.999467 | 0.999453 |
| **101164** | 10 | 72976250 | 72978750 | 10 | 73056250 | 73058750 | 0.998081 | 0.998106 | 0.998049 | 0.998081 | ... | 0.998634 | 0.998489 | 0.995984 | 0.998057 | 0.998444 | 0.998638 | 0.998109 | 0.998574 | 0.998484 | 0.998109 |
| **101165** | 10 | 72971250 | 72973750 | 10 | 73136250 | 73138750 | 0.999703 | 0.999601 | 0.999715 | 0.999709 | ... | 0.999614 | 0.999000 | 0.999752 | 0.999715 | 0.997925 | 0.999616 | 0.999717 | 0.999614 | 0.998971 | 0.999747 |
| **101166** | 10 | 72956250 | 72958750 | 10 | 73136250 | 73138750 | 0.999502 | 0.999943 | 0.999934 | 0.999509 | ... | 0.999978 | 0.999932 | 0.999963 | 0.999489 | 0.999694 | 0.999941 | 0.999506 | 0.999978 | 0.999929 | 0.999513 |
| **101167** | 10 | 72956250 | 72958750 | 10 | 73056250 | 73058750 | 0.998333 | 0.999476 | 0.999779 | 0.998301 | ... | 0.999799 | 0.999760 | 0.999475 | 0.998351 | 0.999483 | 0.999498 | 0.998351 | 0.999799 | 0.999764 | 0.998351 |
| **101168** | 10 | 72951250 | 72953750 | 10 | 72981250 | 72983750 | 0.947861 | 0.949555 | 0.947861 | 0.948079 | ... | 0.948975 | 0.948324 | 0.948755 | 0.947861 | 0.950364 | 0.949129 | 0.947996 | 0.948975 | 0.950512 | 0.948855 |
| **101169** | 10 | 72951250 | 72953750 | 10 | 72976250 | 72978750 | 0.999529 | 0.993542 | 0.999488 | 0.999529 | ... | 0.997486 | 0.994684 | 0.997393 | 0.999535 | 0.994824 | 0.993366 | 0.999529 | 0.995852 | 0.997383 | 0.999529 |
| **101170** | 10 | 73136250 | 73138750 | 10 | 73236250 | 73238750 | 0.953523 | 0.729933 | 0.954445 | 0.952785 | ... | 0.398227 | 0.832376 | 0.830190 | 0.954499 | 0.833134 | 0.953695 | 0.954233 | 0.402025 | 0.831135 | 0.951627 |
| **101171** | 10 | 150831250 | 150833750 | 10 | 150851250 | 150853750 | 0.990458 | 0.995519 | 0.995652 | 0.986731 | ... | 0.995619 | 0.970963 | 0.995505 | 0.998137 | 0.995619 | 0.986625 | 0.993913 | 0.986692 | 0.995529 | 0.997569 |

10 rows × 284 columns

**Table 1:** Overview of Loop Stability for Selected Anchor Pairs Detailed loop stability metrics for the first and last 10 anchor pairs, specified by genomic coordinates (chr1, s1, e1, chr2, s2, e2), from a matrix encompassing 277 maize inbred lines and 101,172 data points.


1. Trieu,T., Martinez-Fundichely,A. and Khurana,E. (2020) DeepMILO: a deep learning approach to predict the impact of non-coding sequence variants on 3D chromatin structure. *Genome Biol.*, 21, 79.

2. Human Genome Overview - Genome Reference Consortium.

3. CSHL Ensembl Plants Zea mays Gene Set.
